# A novel vitamin C analog acts as a potent bio-enhancer to augment the activities of anti-tuberculosis drugs against *Mycobacterium tuberculosis*

**DOI:** 10.1101/2023.11.26.568727

**Authors:** Naveen Prakash Bokolia, Kingsuk Bag, Biplab Sarkar, Ruchi Jhawar, Dipankar Chatterji, Narayanaswamy Jayaraman, Anirban Ghosh

**Affiliations:** Molecular Biophysics Unit, Indian Institute of Science, Bangalore, India; Department of Organic Chemistry, Indian Institute of Science, Bangalore, India

**Keywords:** Mycobacterium tuberculosis, anti-tuberculosis drugs, bio-enhancer agents, kill kinetics, efficacy, antibiotic resistance, reactive oxygen species

## Abstract

*Mycobacterium tuberculosis* is a deadly pathogen that claims millions of lives every year. Current research focuses on finding new anti-tuberculosis drugs that are safe and effective, with lesser side effects and toxicity. One important approach is to identify bio-enhancers that can improve the effectiveness of anti-tuberculosis drugs, resulting in reduced doses and shortened treatment times. We investigated the use of vitamin C-derived isotetrones as bio- enhancer agents. In this context, our results revealed that the lead compound C11 inhibits growth, improves MIC/MBC, and enhances the killing of *M. tuberculosis* H37Rv strain when used in combination with first-line and injectable anti-TB drugs in a dose-dependent manner. The combination of C11 and rifampicin also reduced the generation of spontaneous mutants against rifampicin and reached a mutation prevention concentration (MPC) with moderate rifampicin concentrations. The identified compounds were proven to be effective against the MDR strain of *M. tuberculosis* and non-cytotoxic in HepG2 cells. We also found that C11 induced the generation of reactive oxygen species (ROS) inside macrophages and within bacteria, resulting in better efficacy.

## Introduction

*M. tuberculosis* is the causative agent of tuberculosis disease and has characterized properties that allow it to persist longer within the host organism (1). According to Global Tuberculosis Report 2022, the global TB burden is still high where 66% of patients (non-MDR TB) got treated, whereas only 43% of patients got treated in the case of MDR/RR-TB (2). Several studies have shown that dormant TB bacilli can re-emerge into actively growing pathogens and often give rise to resistant mutants by spontaneous or drug-induced re-activation (3, 4). Due to its persistent nature, the standard TB treatment goes on for a lengthy 6 months often associated with poor compliance and adverse side effects on the patients (5, 6). To counter this multifaceted problem, novel anti-tuberculosis drugs are needed (7, 8), and simultaneously specific bio-enhancer compounds are required to increase the potency of the existing antibiotics in the context of better efficacy, lower concentration and shorter treatment regimens(9). Several studies have demonstrated the uses of bio-enhancers in clinical studies. In one study, the piperine combination reduced the dose level of rifampicin (10). In another study, colistin was shown to increase the bio-efficacy of anti-TB drugs, where colistin helps to increase the permeability of mycobacterial cell walls for anti-TB drugs (11), although colistin is a drug, that is already in use against MDR Gram-negative bacterial infections (12). Bio-enhancers also have been shown to increase the bioavailability of the drug and simultaneously mediate significant advantages to TB patients during the treatment course (13). Importantly, bio- enhancers might reduce the duration and cost of first-line TB drugs which would be a great help to poor patients, government agencies and non-government organizations. Previous studies have shown the anti-tuberculosis activity of vitamin C that potentiates the killing in combination with rifampicin and isoniazid (14, 15). In the present study, we report the effect of vitamin C-derived isotetrones that could significantly increase the activity of TB drugs at *in vitro* and *ex vivo* conditions. Our data further suggested that C11 was able to prevent resistant mutant generation and stimulate ROS production inside cells to attain superior killing. Finally, our compounds proved to be effective against MDR strains of *M. tuberculosis* with a significant drop in MIC of first-line drugs. To summarize, we have identified and validated a C-4 modified isotetrone as a novel class of synthetic bio-enhancer that could have a significant impact on drug regimens for tuberculosis treatment.

## Results

### C-4 modified isotetrone 11 demonstrates significant potentiation of anti-TB drugs

We screened the selected C-4 modified isotetrones (16) **(Figure S1)** for their potential activity against *M. tuberculosis* H37Rv by minimum inhibitory concentration (MIC) assay The compounds were screened alone and also in combination with first-line TB drugs. Simultaneously, the selected C-4 modified isotetrones were analyzed for the planktonic growth inhibitory activity against *M. tuberculosis* H37Rv **(Figure S2)**. These preliminary studies led to the identification of one isotetrone compound 11 (C11) as a potential bio-enhancer agent. The C11 compound exhibited significant MIC modulation of first-line and injectable anti-TB drugs in a dose-dependent manner. In the case of rifampicin, a 66-fold decrease in MIC value was observed (**Table 1**) in combination with C11 (200 μg/ml final concentration). C11 compound has also been shown to improve the efficacy of isoniazid with 16-fold MIC modulation (**Table 1**). Similarly, MIC improvement was also observed in the case of second-line injectable drugs (amikacin and kanamycin) (**Table 1**). Next, we compared the MBC values of the same antibiotics alone or in combination with C11 and we found significant fold changes for rifampicin, isoniazid, kanamycin and amikacin. Specifically, the MBC value of rifampicin reduced to 0.0018 μg/ml (133 fold) in the presence of C11 (200 μg/ml), that combination was able to kill 99.9% of *M. tuberculosis* bacilli (**Table 2**). These results suggested that the presence of compound C11 significantly potentiates the *in vitro* efficacy of anti-TB drugs.

**Table 1.**
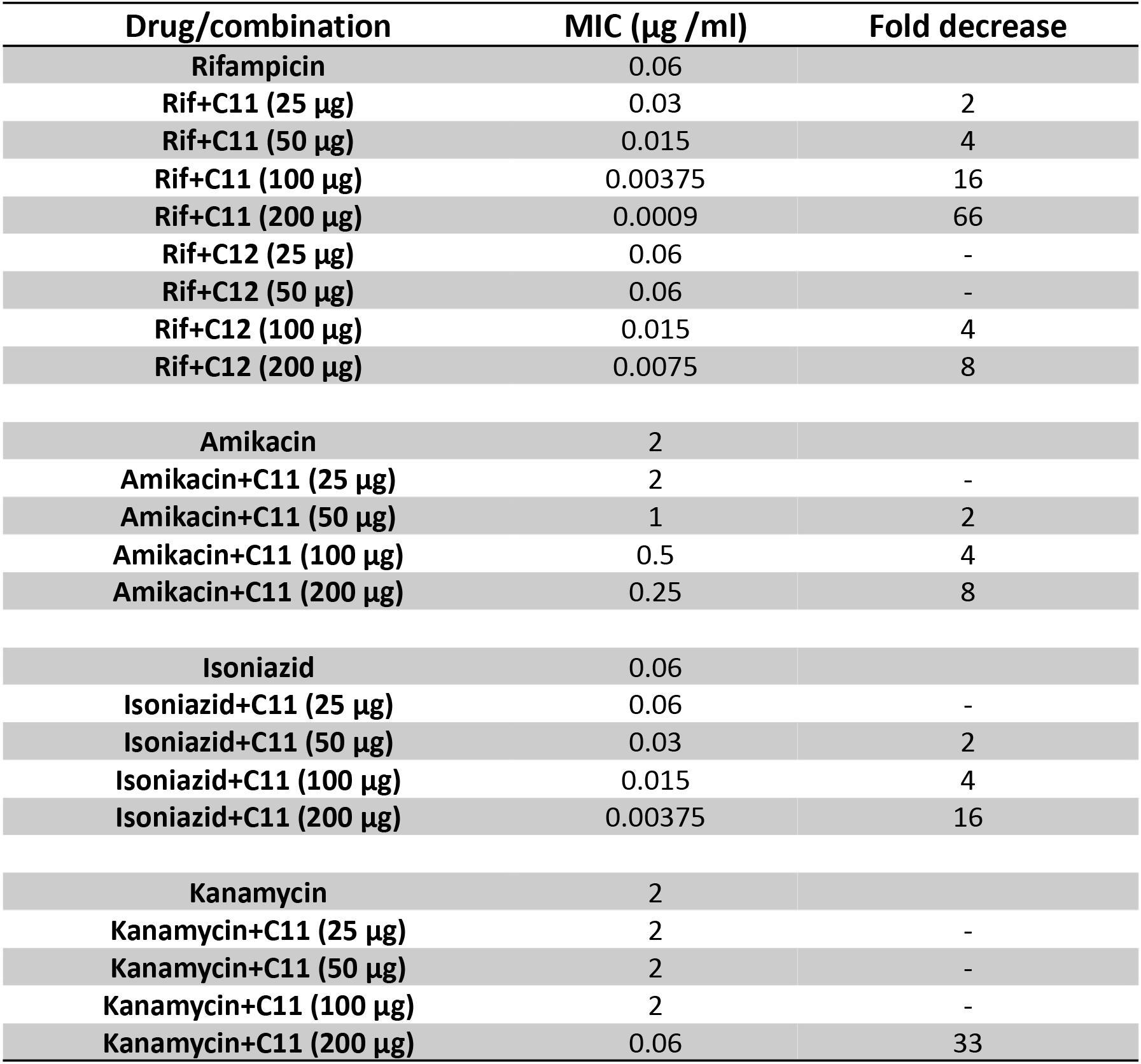
List of MIC values of drug/compound C11 (different concentrations) combinations against *M. tuberculosis* H37Rv. The respective fold decrease in MIC values (compared to the drug alone) is provided in the right column.

| Drug/combination | MIC (µg /ml) | Fold decrease |
| --- | --- | --- |
| Rifampicin | 0.06 |  |
| Rif+C11 (25 µg) | 0.03 | 2 |
| Rif+C11 (50 µg) | 0.015 | 4 |
| Rif+C11 (100 µg) | 0.00375 | 16 |
| Rif+C11 (200 µg) | 0.0009 | 66 |
| Rif+C12 (25 µg) | 0.06 | - |
| Rif+C12 (50 µg) | 0.06 | - |
| Rif+C12 (100 µg) | 0.015 | 4 |
| Rif+C12 (200 µg) | 0.0075 | 8 |
| Amikacin | 2 |  |
| Amikacin+C11 (25 µg) | 2 | - |
| Amikacin+C11 (50 µg) | 1 | 2 |
| Amikacin+C11 (100 µg) | 0.5 | 4 |
| Amikacin+C11 (200 µg) | 0.25 | 8 |
| Isoniazid | 0.06 |  |
| Isoniazid+C11 (25 µg) | 0.06 | - |
| Isoniazid+C11 (50 µg) | 0.03 | 2 |
| Isoniazid+C11 (100 µg) | 0.015 | 4 |
| Isoniazid+C11 (200 µg) | 0.00375 | 16 |
| Kanamycin | 2 |  |
| Kanamycin+C11 (25 µg) | 2 | - |
| Kanamycin+C11 (50 µg) | 2 | - |
| Kanamycin+C11 (100 µg) | 2 | - |
| Kanamycin+C11 (200 µg) | 0.06 | 33 |

**Table 2.** List of MBC values of drug/compound C11 combination against *M. tuberculosis* H37Rv. The respective fold decrease in MBC values (compared to the drug alone) is provided in the right column.

| Drug/combination | MBC (µg /ml) | Fold decrease | MBC/MIC |
| --- | --- | --- | --- |
| Rifampicin | 0.25 |  | 4 |
| Rif+C11 (25 µg) | 0.125 | 2 | 4 |
| Rif+C11 (50 µg) | 0.03 | 8 | 2 |
| Rif+C11 (100 µg) | 0.015 | 16 | 4 |
| Rif+C11 (200 µg) | 0.001875 | 133 | 2 |
| Rif+C12 (200 µg) | 0.125 | 2 | 16 |
| Amikacin | 4 |  | 2 |
| Amikacin+C11 (100 µg) | 4 | - | 8 |
| Amikacin+C11 (200 µg) | 1 | 4 | 4 |
| Isoniazid | 0.25 |  | 4 |
| Isoniazid+C11 (100 µg) | 0.25 | - | 16 |
| Isoniazid+C11 (200 µg) | 0.03 | 8 | 8 |
| Kanamycin | 2 |  | 1 |
| Kanamycin+C11 (100 µg) | 2 | - | 1 |
| Kanamycin+C11 (200 µg) | 0.125 | 16 | 2 |

### The *in vitro* resistance profile of rifampicin was improved with C11

The impact of the C11 compound was analyzed by the emergence of resistant mutants against rifampicin. *In vitro*, spontaneous mutants against rifampicin (5X MIC) arose at a frequency of 1.32 X 10^-8^, whereas in combination with C11 (200 μg/ml), it became ∼10 fold less i.e. 4.3 X 10^-9^ (**Table 3**). The tiny size and late emergence of colonies were observed on 7H11 agar plates carrying both rifampicin and C11, which further demonstrates the better efficacy of rifampicin in combination with C11. Similar trends have been observed in a higher concentration of rifampicin (10X and 20X MIC) as no resistant mutants could be isolated in combination with compound 11, for up to 6 weeks suggesting that it would be the mutation prevention concentration (MPC) of rifampicin in combination with C11. Whereas resistant mutants arose at a frequency of 4.5 X 10^-9^ and 5.2 X 10^-9^ in the presence of 10X and 20X MIC of rifampicin without co-administration of C11. The tiny size and late emergence of colonies were observed on 7H11 agar plates carrying both rifampicin and C11, which further demonstrates the better efficacy of rifampicin in combination with C11. These results evinced the effect of C11 to suppress the generation of resistant mutants of *M. tuberculosis*.

**Table 3.** Spontaneous mutation frequency values of rifampicin alone and with compound C11 for *M. tuberculosis* H37Rv.

| Drug and compound | Resistance frequency |
| --- | --- |
| Rifampicin 5X | 1.32 X 10<sup>-8</sup> |
| Rifampicin 5X + C11 (200 ug/ml.) | 8.9 X 10<sup>-9</sup> |
| Rifampicin 10X | 4.5 X 10<sup>-9</sup> |
| Rifampicin 10X + C11 (200 ug/ml) | no colonies |
| Rifampicin 20X | 5.2 X 10<sup>-9</sup> |
| Rifampicin 20X + C11 (200 ug/ml.) | no colonies |

### C11 demonstrated potent bactericidal activity in combination with anti-TB drugs

To evaluate the killing efficacy of the C11 compound (as a bio-enhancer) in combination with drugs, the time-kill kinetics experiments were performed. Taking the MIC and MBC data into consideration, we used less concentration (2X MIC concentration) of the drugs, and in all the cases, we saw that the presence of compound C11 manifested a better killing. Specifically, rifampicin (**Figure 1A**), isoniazid (**Figure 1B**), amikacin (**Figure 1C**) and kanamycin (**Figure 1D**) in combination with compound 11 showed an additional 2-3 log kill at day-21. In all cases, a better kill was observed with C11 in a dose-dependent manner where maximum killing was achieved at 200 μg/ml. The kanamycin+C11 (200 μg/ml) combination resulted in complete sterilization of the culture as no CFU was detected at day 21 (**Figure 1C**). When no compounds were present, long-term killing seemed to be less efficient by the drugs, which was evidenced by the re-emergence of resistant mutants (as they were growing in the presence of the drug) at later time points due to low concentration (2X) of the drugs used in the assay **(Figure 1A, 1B, 1C & 1D**). Thus, our data suggested that C11 in combination with existing drugs manifests a better dose-dependent killing in comparison to drug-alone treated cultures and significantly lowers the concentration of existing antibiotics from conventional 10X **(Figure S3)** to 2X MIC concentration and could achieve the same ∼99.9% killing by 9 days and critically suppress the enrichment of mutants in the culture in later time points.

**Figure 1.**
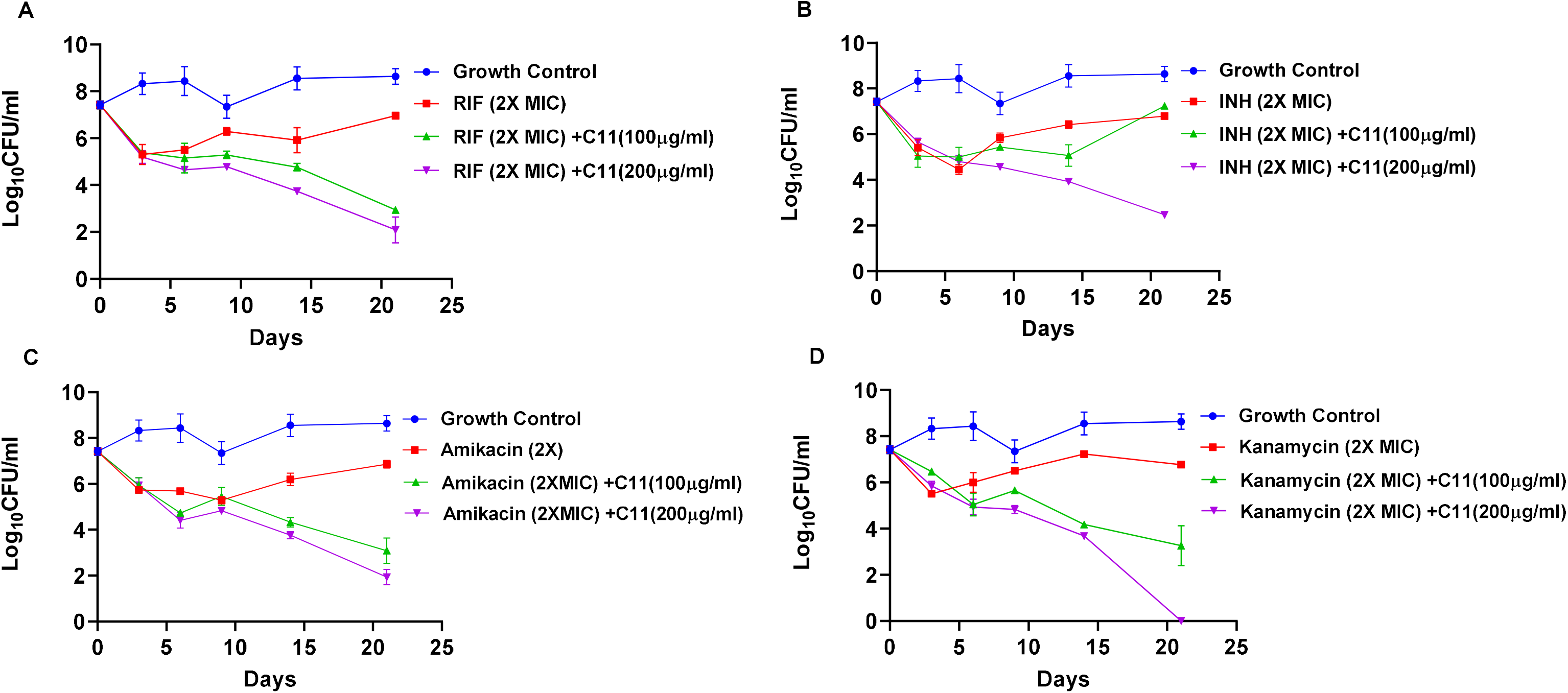
Killing kinetics of (A) Rifampicin, (B) Isoniazid, (C) Amikacin and (D) Kanamycin at 2X MIC concentrations with and without C11 (100 µg/ml and 200µg/ml) against *M. tuberculosis*.

### Rifampicin and isoniazid showed enhanced intracellular killing in combination with C11

The intracellular killing efficacy study was performed in *ex vivo* conditions using cultured murine macrophage cells (RAW 264.7 cell line). The combination of C11 (used at a concentration with no macrophage cytotoxicity i.e. 200 µg/ml, confirmed by cellular cytotoxicity assay) with rifampicin (10X MIC concentration) and isoniazid (10X MIC concentration) was found to be more effective in killing of the *M. tuberculosis* bacilli inside the macrophages. Specifically, the data suggests that C11 with rifampicin showed superior activity in a time-dependent manner, where ∼1.5 log CFU reduction was observed at day 6, which was ∼0.6 log higher than the rifampicin control **(Figure 2A)**. Similar results were also observed in the case of the isoniazid + C11 combination where it showed superior efficacy (∼0.4 log more killing in day 6) compared to isoniazid **(Figure 2B).** These results reasserted the role of C11 as a bio-enhancer and suggested that it might trigger some additional pathways inside the macrophages resulting in a greater killing of the bacteria.

**Figure 2.**
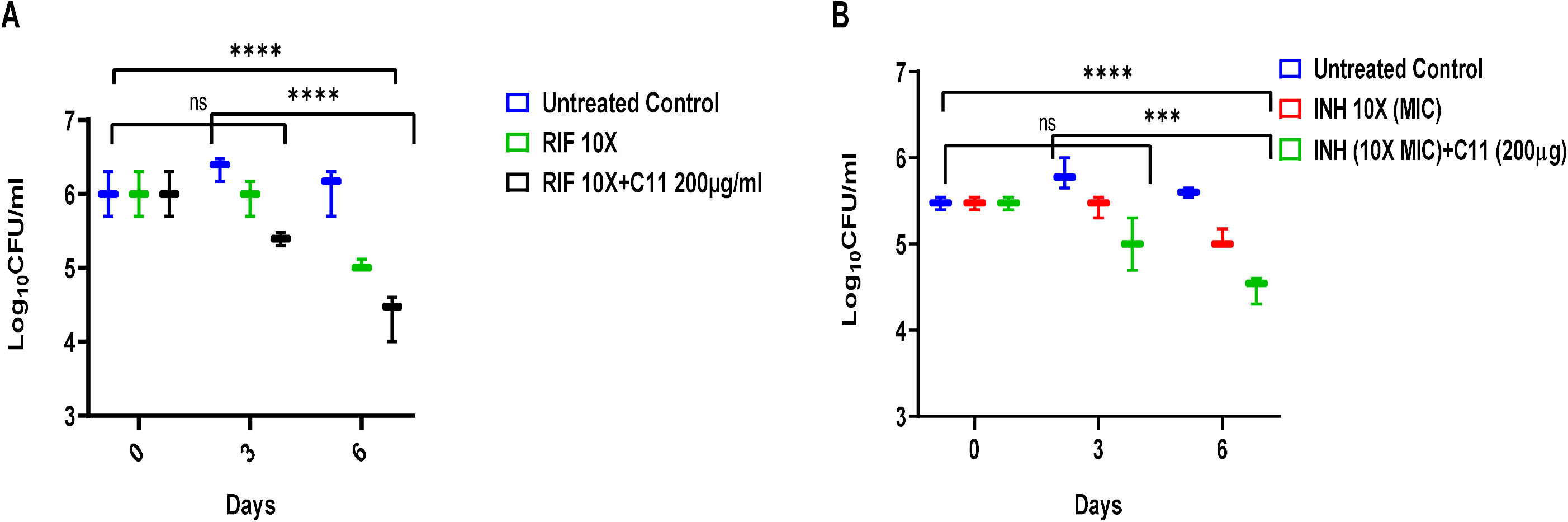
Killing kinetics of (A) rifampicin and (B) Isoniazid with and without C11 in RAW264.7 macrophages infected with *M. tuberculosis*. Two-way ANOVA (multiple comparison test) was used to calculate the statistically significant values: \*\*\*\**p* < 0.0001; \*\*\**p*=0.0001; \**p*=0.05; ns= non-significant.

### Compound C11 demonstrates pro-oxidant properties in macrophage cells

As both the compounds are synthetic analogs of vitamin C, therefore, we first evaluated the properties of compounds as pro-oxidants within mammalian cell lines. *M. tuberculosis* primarily resides within alveolar macrophages thus we examined the effect of compound C11 on murine macrophages. Murine macrophage cells (RAW 264.7) were treated with varying concentrations of C11 and incubated for 24 hours. After incubation, cell-permeable 2’,7’- dichlorodihydrofluorescein diacetate (H2DCFDA) dye was added and fluorescence intensity was determined, which is a direct measurement of the reactive oxygen species (ROS) inside macrophages. The increase in fluorescence was observed in the presence of C11 (in a range of 50 µg/ml to 400 µg/ml) **(Figure 3)** compared to the untreated series highlighting its prooxidant role in stimulating ROS production inside macrophages in a concentration-dependent manner.

**Figure 3.**
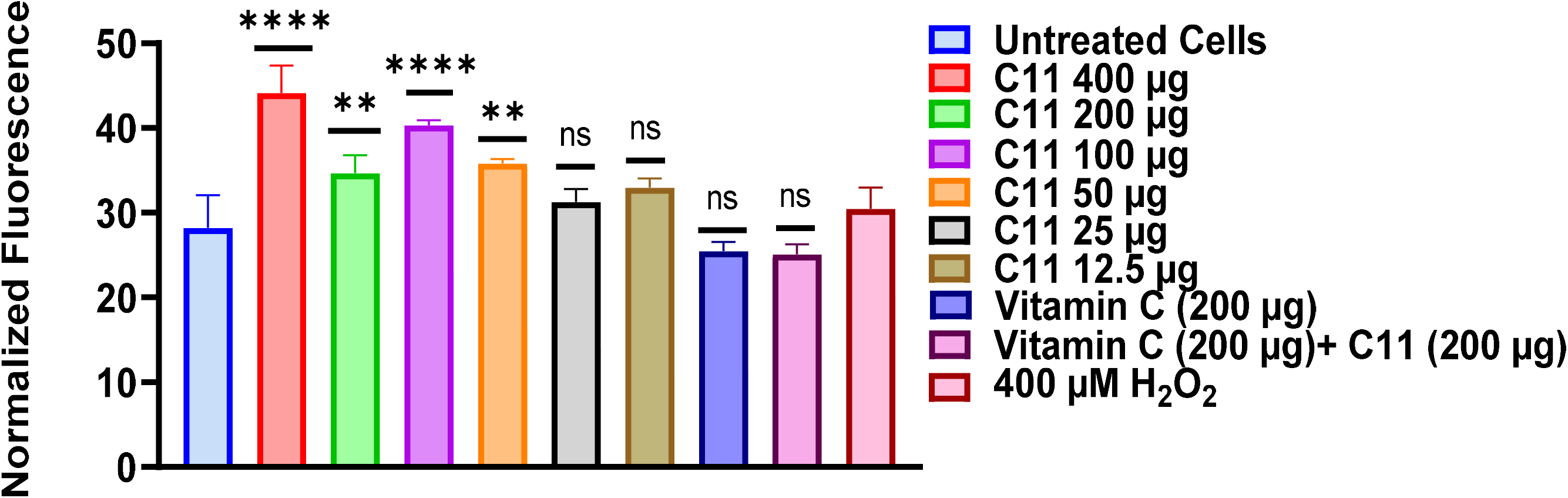
C11 induces ROS species generation inside macrophages (RAW264.7) in a dose-dependent manner. The plot presents the normalized fluorescence values from the H2DCFDA dye treatment. One-way ANOVA (multiple comparisons with respect to control) was used to mention the significant values as follows: 400µg: \*\*\*\**p* < 0.0001; 200µg: \*\**p* = 0.0079; 100µg: \*\*\*\**p* < 0.0001; 50µg: \*\**p*=0.0018; ns= non-significant.

### Compound C11 mediates the direct oxidation of mycothiol and modulates the intracellular redox state of *M. tuberculosis*

To determine the intracellular oxidation level of *M. tuberculosis* cells in response to C11, we used the *M. tuberculosis* Mrx1-roGFP2 strain (mycothiol-mycoredoxin redox-based biosensor strain). This biosensor strain was constructed to sense mycothiol’s oxidation level, and the respective signal was monitored via redox-sensitive ro-GFP2 (13). Mycothiol plays a crucial role in recycling the reduced form of mycoredoxin and the underlying principle is that if any compound induces ROS production, then it will keep the mycothiol in an oxidized state and eventually will shift the cytoplasmic equilibrium towards the oxidized form of mycoredoxin. We treated the mid-log phase culture of *M. tuberculosis* Mrx-roGFP2 strain under different conditions of C11 compound/anti-TB drug combinations for 48 hours. The results were represented by the percentage of heterogeneity in the population for the intramycobacterium redox state. In our first observation, we found that the low concentration of anti-TB drugs (1X MIC) alone helped *M. tuberculosis* cells to encounter reductive stress. This led to a significant shift in subpopulation towards the reduced state after 48 hours **(Figure 4A)**. On the other hand, treatment with compound C11 caused a significant shift in intramycobacterial redox potential towards the oxidized state **(Figure 4A)**. We noticed that a concentration of 200 µg/ml of compound C11 produced a significant effect in generating oxidized subpopulations (about 7.4%). A similar result was observed when culture was treated with a combination of C11 and different anti-TB drugs. This feature highlights an important mechanism of C11 as a bio- enhancer agent in the enhanced killing of *M. tuberculosis* cells with low concentrations of anti-TB drugs that remain persistent even after 48 hours. Furthermore, we found that the C11- treated “oxidized subpopulation” was not neutralized or suppressed with thiourea treatment, whereas the H_2_O_2_-mediated “oxidized subpopulation” was significantly neutralized (reduced or basal) with thiourea **(Figure 4B)**. Additionally, we observed that the treatment of vitamin C (at 1mM or 10mM) alone or in combination with rifampicin caused reductive stress **(Figure 4A)**. Previous studies have also reported that vitamin C can induce reductive stress in THP-1 cells infected with *M. tuberculosis* Mrx-1 roGFP2 strain (15), but our findings suggested that the combination of vitamin C and rifampicin led to a greater shift towards the reduced (mycothiol) subpopulations. This could be due to the low drug concentration and longer treatment duration (48 hours) that was selected for our experiment, which was to gain a better understanding of the properties of the C11 compound in targeting *M. tuberculosis*. In summary, our results demonstrate that C11 plays a significant role in the oxidation of mycothiol and disruption of cytoplasmic redox homeostasis.

**Figure 4.**
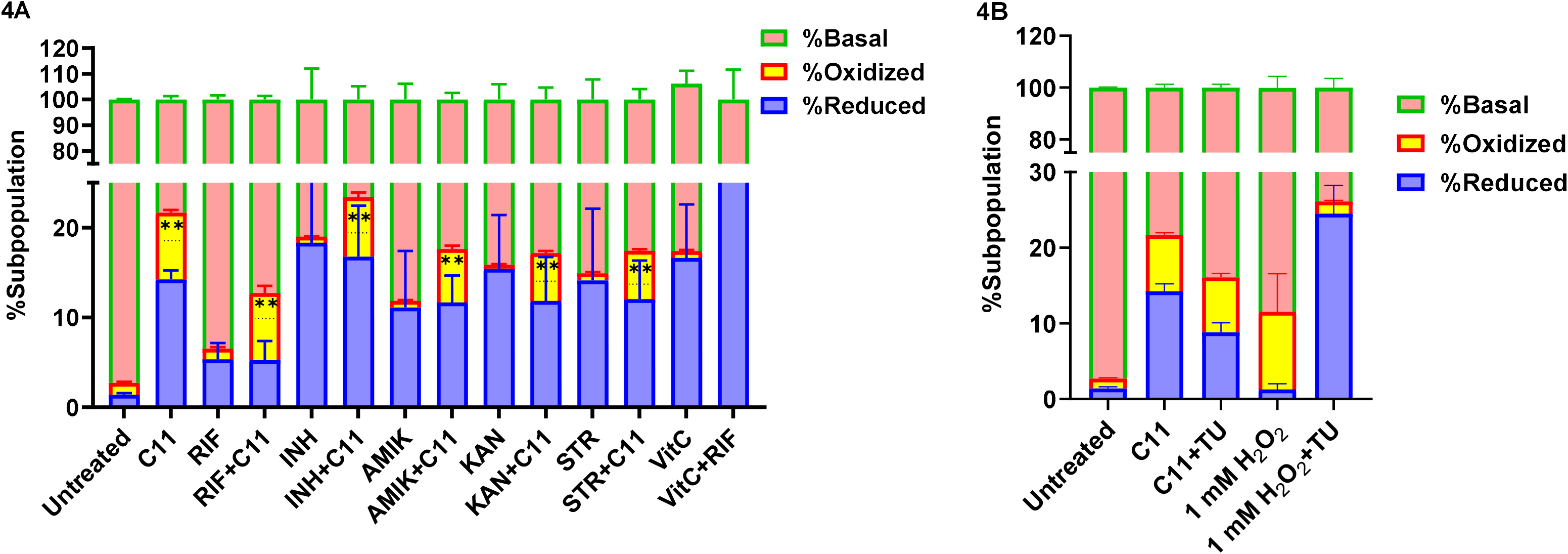
(A) C11 compound mediates direct oxidation of mycothiol inside *M. tuberculosis*. The plot represents the redox status of the Mrx1-roGFP2 strain in response to C11 alone and drug combination. The %subpopulation (oxidized, reduced and basal) of Mrx1-roGFP2 strain was determined after 48 hours of treatment (incubation), and data were plotted. **(B)** The plot presents the redox status of cells treated with C11 and H_2_O_2_ in the presence and absence of thiourea (TU). The plot was analyzed by using “Sidak’s multiple comparisons test” “%Oxidized vs. %reduced” and significant values are as follows \*\**p*=0.0023; \*\*\*\**p*<0.0001.

### C11 sensitizes the MDR strain of *M. tuberculosis* to rifampicin, isoniazid and streptomycin

Based on the positive findings in *M. tuberculosis* H37Rv (wildtype strain), we determined the efficacy of the C11 and anti-TB drugs combinations against the clinically isolated multidrug-resistant strain of *M. tuberculosis* (NHN1664; catalog No. NR-19016), that was originally isolated in China and obtained through BEI Resources, NIAID, NIH. This MDR strain is resistant to rifampicin, isoniazid and streptomycin. Strikingly, the MIC results showed that the MDR strain became sensitive to rifampicin (MIC: 0.25 µg/ml) in the presence of C11 (200 µg/ml final concentration), whereas the rifampicin alone had a high MIC value >128 µg/ml, indicating a fold change of >512 **(Table 4)**. Significant modulation of MIC values was also observed for isoniazid (MIC fold change: 16) and streptomycin (MIC fold change: >1024) **(Table 4)**. However, no such fold change in MIC value was observed for moxifloxacin, like in *M. tuberculosis* H37Rv. It is important to note that the tested concentration of C11 for all assays (200 µg/ml) was not able to suppress the growth of the bacteria in MIC plates by itself. In the presence of C11, the MBC values of rifampicin, isoniazid and streptomycin were also improved by >128-fold, 64-fold, and >128-fold respectively **(Table 5)**. Altogether, these results suggested that compound C11 could cause a reversal of resistance and make cells vulnerable to the same existing TB drugs against which they developed resistance before.

**Table 4.** List of MIC values of drug/compound C11 (200µg/ml fixed concentration) combinations against *M. tuberculosis* NHN1664 (multi-drug resistant; MDR) strain. The respective fold decreases in MIC values (compared to the drug alone) are provided in parallel.

| Drug/combination | MIC (µg /ml) | Fold decrease |
| --- | --- | --- |
| Rifampicin | >128 µg/ml |  |
| Streptomycin | >256 µg/ml |  |
| Isoniazid | 16 µg/ml |  |
| Moxifloxacin | 0.6 µg/ml |  |
| C11 (200 µg/ml) | Inactive |  |
| Vitamin C (200 µg/ml) | Inactive |  |
| Rifampicin+C11 | 0.25 µg/ml | >512 |
| Streptomycin+C11 | 0.25 µg/ml | >1024 |
| Isoniazid+C11 | 1 µg/ml | 16 |
| Moxifloxacin+C11 | 0.6 µg/ml | No Change |
| Rifampicin+ Vitamin C (200 µg/ml) | >128 µg/ml | No Change |

**Table 5.** List of MBC values of drug/compound C11 (200µg/ml fixed concentration) combinations against *M. tuberculosis* NHN1664 (multi-drug resistant; MDR) strain. The respective fold decreases in MIC values (compared to the drug alone) are provided in parallel.

| Drug/combination | MBC (µg /ml) | Fold decrease |
| --- | --- | --- |
| Rifampicin | >256 µg/ml |  |
| Streptomycin | >256 µg/ml |  |
| Isoniazid | 128 µg/ml |  |
| Moxifloxacin | 0.25 µg/ml |  |
| Rifampicin+C11 | 2 µg/ml | >128 |
| Isoniazid+C11 | 4 µg/ml | 64 |
| Streptomycin+C11 | 2 µg/ml | >128 |
| Moxifloxacin+C11 | 0.25 µg/ml | - |

### Compounds C11 demonstrated a high safety index (SI) profile

Finally, we examined the safety profile of the C11 compound by MTT assay on human liver cancer cell line HepG2, where C11 was found to be significantly safe up to 2000 µg/ml **(Figure 5)**. C11 exhibited higher CC50 values, where no significant reduction of the live cells (metabolic activity) was observed at 250 µg/ml concentration (∼94% live cells).

**Figure 5.**
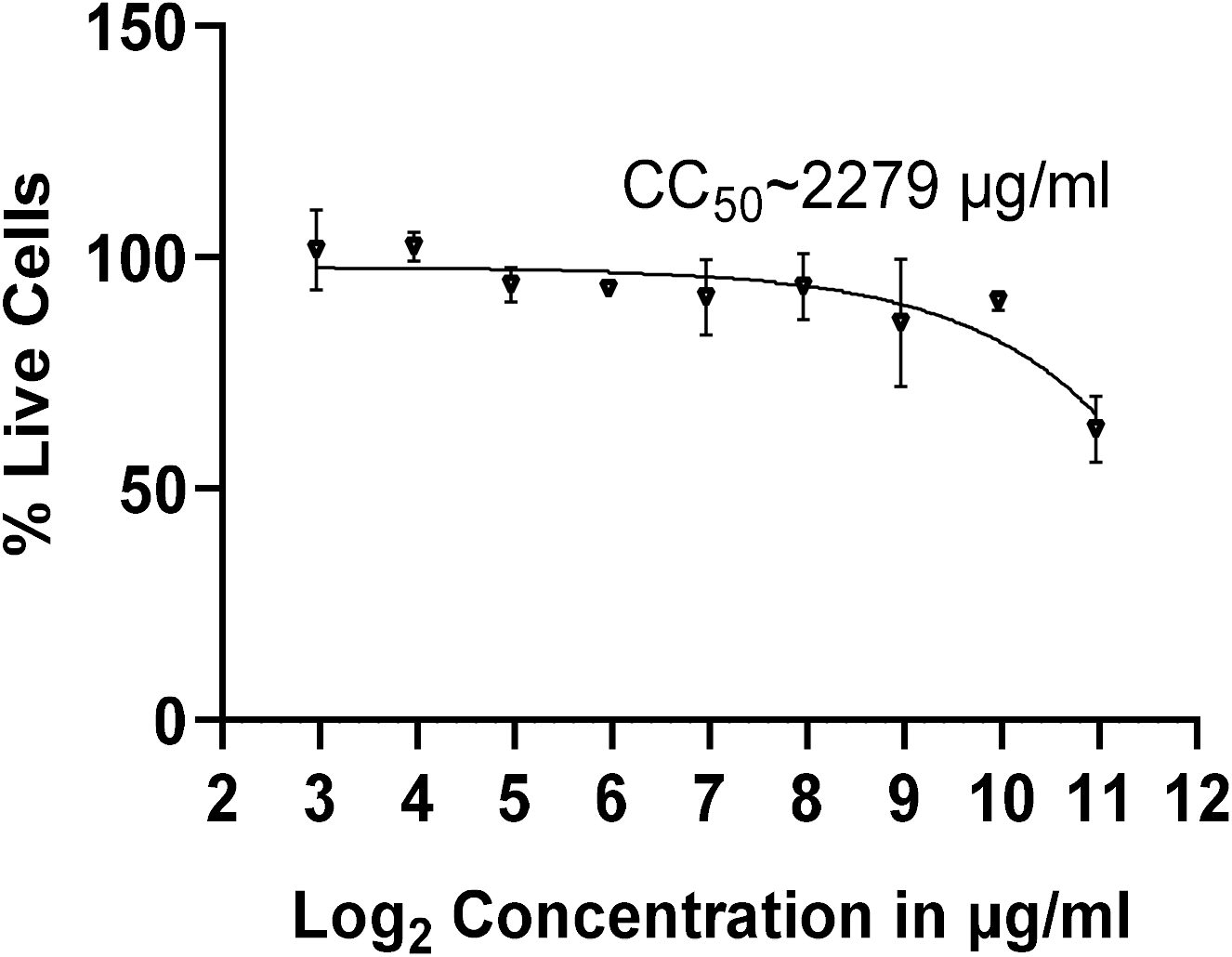
The plot presents the CC_50_ value of the C11 compound in the HepG2 cell lines. The graph is plotted between the %live cells vs log_2_ (concentration) of the compound C11. The CC_50_ value was determined by the non-linear fit method in GraphPad Prism and further selected the [inhibitor] vs. response (three parameters), and a total of 9 points from the Y-axis were analyzed.

## Discussion

In the present study, we identified the specific bio-enhancer compound that could improve the bactericidal activity of existing TB drugs. As the compound C11 was synthesized by attaching L-phenylalanine to the vitamin C core structure, those could also be regarded as vitamin C analogs (16). The identified bio-enhancer C11 has been shown to have a potent effect in combination with first-line and second-line injectable drugs. The identified bio-enhancer follows Lipinski’s rule that involves: molecular weight <500 daltons; <5 hydrogen bond donors; <10 hydrogen bond acceptors & octanol-water partition coefficient log P value <5 (17). Our findings indicate that when C11 is used in conjunction with different anti-TB drugs, it results in a more effective dose-dependent elimination of *M. tuberculosis* under *in vitro* and *ex vivo* conditions and importantly reduces the concentration of antibiotics from 10 times to just 2 times of MIC. Our study also showed that bio-enhancer C11 can potentially help to suppress the resistant mutant enrichment, reach a mutation prevention concentration (MPC) of rifampicin and also make *M. tuberculosis* MDR strain susceptible to both rifampicin and isoniazid. This could be a significant breakthrough in dealing with resistant mutants and could help preserve second-line drugs from overuse. A previous study showed that vitamin C treatment resulted in the production of free hydroxyl radicals in *M. tuberculosis*, through Fenton’s reaction (18). This same study also showed that vitamin C tends to increase the ferrous ion levels within *M. tuberculosis* bacilli. In this perspective, the C11 (an analog of vitamin C) could induce a similar Fenton’s reaction, and maintain better bactericidal activity when combined with anti-TB drugs (19, 20). It’s critical to note that our research demonstrated the potential of C11 in lowering the MIC of rifampicin (and other drugs) in the MDR strain. The results indicated that in the presence of C11 and antibiotics (rifampicin, isoniazid, and streptomycin), the level of ROS species increased that includes superoxide or hydroxyl radical formation by either Fenton’s reaction or in mediating the direct oxidation of NADH. This increase in ROS species might be able to restore sensitivity to the respective antibiotic in the mutant strain. Our research demonstrated evidence of pro-oxidant property of C11 by the oxidation status of mycothiol (in the Mrx1-roGFP2 strain). Firstly, we noted that low drug concentration (1XMIC) allowed the bacterial population to maintain the reduced state of mycothiol, where the dissipation of the reduced state is an important mechanism for the adaptation of mycobacteria against oxidative stress (21). Simultaneously, we also observed an increased intra-macrophage and intra-mycobacterium oxidation level in combination with drugs (and C11 alone), which persisted even after 48 hours of treatment. Importantly, the C11- mediated mycothiol oxidation was not neutralized/suppressed by the thiourea treatment, unlike the H_2_O_2_ treatment. This observation points toward the production of ROS species upon C11 treatment, which induced multiple oxidative stress (importantly oxidation of cytoplasmic mycothiol) responses inside *M. tuberculosis* resulting in increased susceptibility to anti-TB drugs. Finally, compound C11 was found to be non-cytotoxic and effective in maintaining the better intracellular oxidative environment/metabolic activity of the HepG2 cells determined by MTT assay (indirect correlation with NADH and NADPH levels). In summary, this study demonstrated that the novel vitamin C analog C11 can enhance bioactivity against WT and MDR strains of *M. tuberculosis* under *in vitro* and *ex vivo* conditions and importantly reduce the likelihood of generating resistant mutants to the first-line TB drug, rifampicin, resulting in a superior therapeutic outcome. Further studies are needed to develop orally active formulations with rifampicin and isoniazid and validate bioavailability and efficacy on *M. tuberculosis* infected animal models along with first-line TB drugs.

## Materials and methods

### Bacterial strains, media and growth conditions

1. *M. tuberculosis* H37Rv strain was grown and maintained in 7H9 medium supplemented with 10% oleic acid-albumin-dextrose-catalase, 0.05% (vol/vol) Tween 80 (Sigma, St. Louis, MO), and 0.2% (vol/vol) glycerol or on Middlebrook 7H11 agar (7H11) supplemented with 10% (vol/vol) oleic acid-albumin-dextrose-catalase (Difco) (22). Cultures were grown and maintained in roller bottles (in 7H9 medium) or 7H11 agar plates. The MDR strain NHN1664 (NHN 1664, catalog No. NR-19016) was used in this study, originally isolated in China and obtained through BEI Resources, NIAID, NIH. This MDR strain of *M. tuberculosis* exhibits resistance to first line drugs: rifampicin, isoniazid, ethambutol including streptomycin (aminoglycoside antibiotic).

### Compounds/antibiotics used in this study

The bio-enhancer agent isotetrone 11 (C11) was dissolved in DMSO (20 mg/ml for MIC; 40mg/ml for HepG2 and RAW264.7 cell line studies), and used at mentioned concentrations in MIC and kill kinetics experiments (DMSO concentration remained below 1% in experimental plates/tubes). Rifampicin was dissolved in DMSO. Amikacin, kanamycin, isoniazid and streptomycin were dissolved in water and used for described experiments.

### Minimum Inhibitory Concentration (REMA) assay

MIC experiments were performed by broth dilution (two-fold serial dilution) method against *M. tuberculosis* H37Rv and MDR strain of *M. tuberculosis* (NHN 1664) (23). Firstly, the drug alone was serially diluted (in 7H9 medium without glycerol) in a horizontal manner (columns 1-10 of 96 well plates), followed by compounds that were added at a fixed concentration in a particular row and varied from one row to another that gives 200µg/ml, 100 µg/ml, 50 µg/ml and 25 µg/ml of compound concentrations. The *M. tuberculosis* culture was adjusted at 10^6^ CFU/ml and added into each well. MIC plates were incubated for 7 days at 37°C. Resazurin dye was added after the 7^th^ day, and plates were incubated further at 37°C for 24 hours. The MIC was determined as the minimum antibiotic concentration at which the resazurin dye did not change to pink.

### Minimum Bactericidal Concentration determination

As described earlier (24), from MIC plate wells, where no growth was observed 100 µl of cells were spread on 7H11 agar plates and CFU was estimated after 4 weeks at 37°C. MBC values were determined as the minimum concentration of the drug where 99.9% killing of *M. tuberculosis* bacilli was achieved.

### Estimation of resistance frequency

The single-step method was used to isolate spontaneous resistant mutants of *M. tuberculosis* against rifampicin. As previously described (25), *M. tuberculosis* H37RV strain was grown to a mid-exponential phase with an optical density (OD_600_) of 0.6 and the culture was then centrifuged and concentrated 100 times to attain a bacterial count of 10^10^/ml. 100 μl of the culture was plated in triplicate on MB7H11 agar plates containing rifampicin and the remaining cultures were diluted till 10^-8^ dilution and spread on drug-free MB7H11 agar plates for CFU determination (drug-free control). Mutation frequencies were calculated by dividing the number of colonies on a rifampicin-containing plate by the number of colonies on the drug-free plate after 4-6 weeks. Resistant mutant colonies were further streaked on MB7H11 +rifampicin plates to confirm rifampicin resistance and a few colonies were taken for further characterization.

### *In vitro* time kill assay

As previously described (26), the *M. tuberculosis* culture was grown and harvested when it reached mid exponential phase. The density of the culture was adjusted at OD∼0.5 which gives 10^8^ CFU/ml and was determined along with the experiment. The inoculum was diluted 1/10 times, which gives ∼10^7^ CFU/ml and the resultant diluted culture was dispensed in experimental culture tubes. The compounds were added in combination with first-line drugs in replicates (experimental culture tubes). Experimental culture tubes were incubated at 37°C under shaking conditions. The culture aliquots were removed at different time points (in days) for CFU enumeration. The culture aliquots were washed/centrifuged with NST (normal saline tween) to remove residual drugs and compounds. After washing, 10-fold serial dilution was performed up to 10^-6^ dilution, and the aliquots from the respective dilution were plated on 7H11 agar medium for CFU enumeration. The CFU was counted after 14-16 days and the graph was plotted using Log_10_ CFU/ml (y-axis) vs time (x-axis).

### Intracellular kill kinetics assay

The Raw264.7 cell line was used to determine the potency of compounds against intracellular *M. tuberculosis* within the macrophages (27). The cell line was grown and maintained in DMEM (Dulbecco’s Modified Eagle Medium) complete medium. The cell lines were seeded in 24 well plates with 5x10^5^ cells. Cells were infected with 1x10^7^ CFU/ml at an MOI (multiplicity of infection) of 100 and incubated for 4 hours at 37°C with 5% CO_2_. After infection, macrophages were treated with DMEM-negative media (without antibiotic) containing only amikacin (100 µg/ml) for 10 minutes and washed thrice with phosphate buffer saline (PBS) to eliminate any extracellular bacteria. The desired concentrations of compounds in combination with drugs prepared in DMEM (without antibacterial drugs) were added into the experimental wells (in triplicates). Infected macrophages were lysed by 0.25% SDS in phosphate buffer saline, followed by serially diluted in PBS, and aliquots were spotted on 7H11 agar plates for CFU enumeration.

### Cellular cytotoxicity determination by MTT [3-(4, 5-dimethylthiazol-2-yl)-2, 5- diphenyltetrazolium bromide] assay

Human HepG2 cell lines were seeded and maintained in DMEM (catalog number: 10566016) complete media. Cells were seeded with a density of 1x10^5^ cells/well. Upon the attachment of cells, the compound concentrations were serially diluted in DMEM complete medium and added into each well (in triplicates of each concentration). Tamoxifen (cytotoxic concentrations) was used as a positive control for HepG2 cells. After the treatment of compounds, the 96-well plate was incubated for 24 hours at 37°C, 5% CO_2_. After, 24 hours, compounds containing media were removed and fresh media was added, subsequently, MTT dye was added and incubated further for 3 hours. The resultant purple formazan was dissolved in DMSO, and absorbance was determined at 570 nm.

### Determination of pro-oxidant properties of the compounds in macrophage cells

The pro-oxidant properties of the compounds were determined in RAW264.7 (murine macrophage) cell lines. The cell lines were seeded with a density of ∼1x10^5^ cells/well. Each compound alone and in combination were serially diluted in the DMEM complete medium that yielded a range of concentrations and added into the respective wells (in triplicates of each concentration). After the treatment of compounds, the 96-well plate was incubated for 24 hours at 37°C, 5% CO_2_. The 400µM H_2_O_2_ was used as the positive control, and cells were treated for 1 hour (28). After incubation, media was removed and wells were washed with PBS, followed by DMEM complete media (without phenol red) and H2DCFDA dye was added at the final concentration of 20µM. The H2DCFDA dye-treated plate was further incubated for 20 minutes (time to allow the dye permeability through the plasma membrane). The fluorescence was measured with the excitation and emission at 495 and 527nm respectively, and readings were subtracted from the background (H2DCFDA dye only).

### Determination of intramycobacterial reactive oxygen species (*E_msh_*)

To determine the ROS level within *M. tuberculosis*, we used the redox-sensitive strain Mrx1- roGFP2, that is the mycothiol (MSH)-mycoredoxin based biosensor strain (29). Mycothiol is a unique compound that efficiently maintains the reduced state within the cytoplasm of mycobacteria and recycles the oxidized mycoredoxin into a reduced form. The underlying basis of using Mrx1-roGFP2 strain was to determine the ROS property of bio-enhancer compound C11, which could oxidize the mycothiol and subsequently respective emission signal is detected from the GFP. Upon oxidation of mycothiol, the emission signal at 510 nm is analyzed, where excitation at 405 nm the emission is higher whereas the excitation at 488 nm the emission signal is lowered. The opposite (lowered emission with excitation at 405 nm and higher emission at 488 nm) is in the case of reduction of mycothiol. The biosensor strain was grown in 7H9 medium supplemented with 10% ADC and hygromycin (50 µg/ml) and grown till the mid-exponential phase (O.D.∼0.6). At this point, the culture was harvested and density was adjusted at 10^7^ CFU/ml. The culture was treated in PBS with different conditions in triplicates and further incubated for 48 hours. After incubation, samples were subjected to data acquisition by using BD FACS ARIA FUSION. The population distribution (basal, reduced and oxidized) of *M. tuberculosis* and data analysis were performed by using BD FACSDiva software v 8.0.3.

## Acknowledgment

We are grateful to Dr. Amit Singh (Centre for Infectious Disease Research, Indian Institute of Science, Bangalore) for kindly providing the *M. tuberculosis* NHN1664 (MDR) strain and *M. tuberculosis* Mrx1-ro-GFP2 strain. We acknowledge with Mrs. Madhumol Jeevan, Indian Institute of Science, Bangalore for her help in acquisition and analysis if of FACS data of *M. tuberculosis* samples.

## Author contributions

D.C., N.J. and A.G. conceived the study. N.B., K.B. and A.G. designed the research; N.B., K.B., B.S. and R.J. performed experiments; N.B., N.J., D.C. and A.G. analysed the data; N.B., N.J. and A.G. wrote the paper.

List of Supplementary Materials: Figures S1, S2 and S3.

