## Supplementary data for "A novel vitamin C analog acts as a potent bio-enhancer to augment the activities of anti-tuberculosis drugs against *Mycobacterium tuberculosis*"

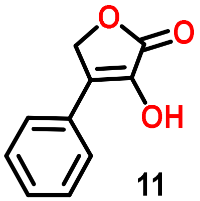

11

Chemical Formula: $C_{10}H_8O_3$
Exact Mass: 176.0473

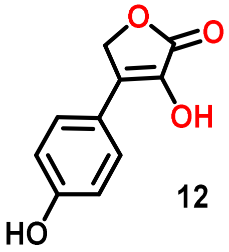

12

Chemical Formula: $C_{10}H_8O_4$
Exact Mass: 192.0423

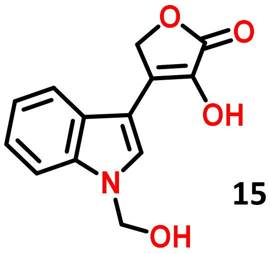

15

Chemical Formula: $C_{13}H_{11}NO_4$
Exact Mass: 245.0688

**Figure S1:** Structure of C-4 modified isotetrones (C-11, C12 and C-15) those were screened (in combination studies) against *M. tuberculosis* H37Rv.

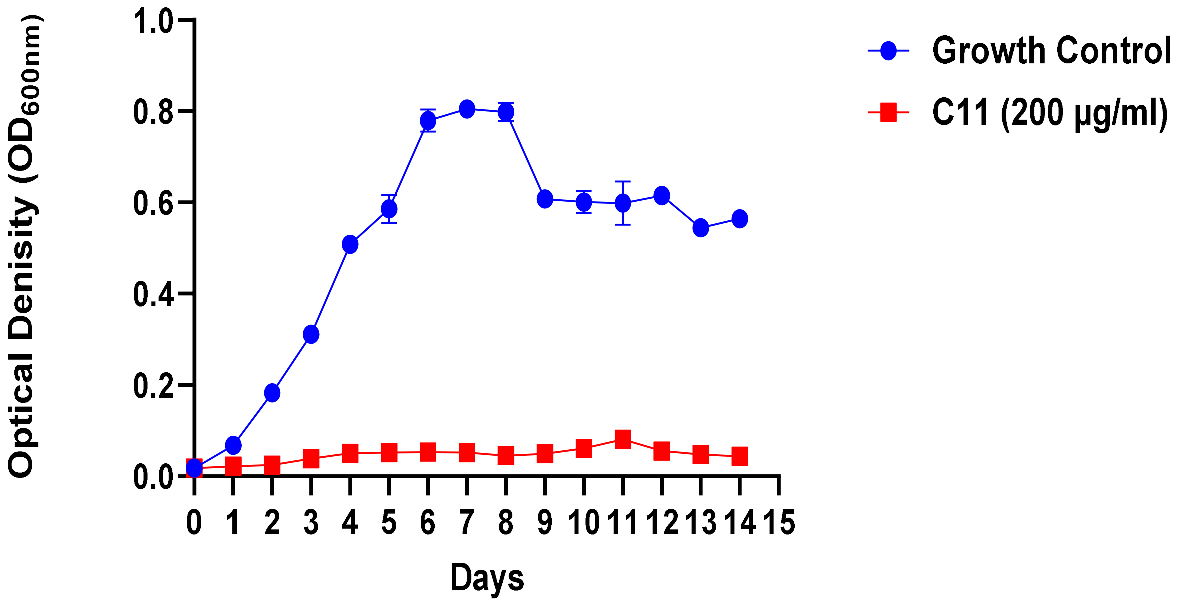


**Figure S2:** C11 isotetrone interferes with the planktonic growth of *M. tuberculosis*.

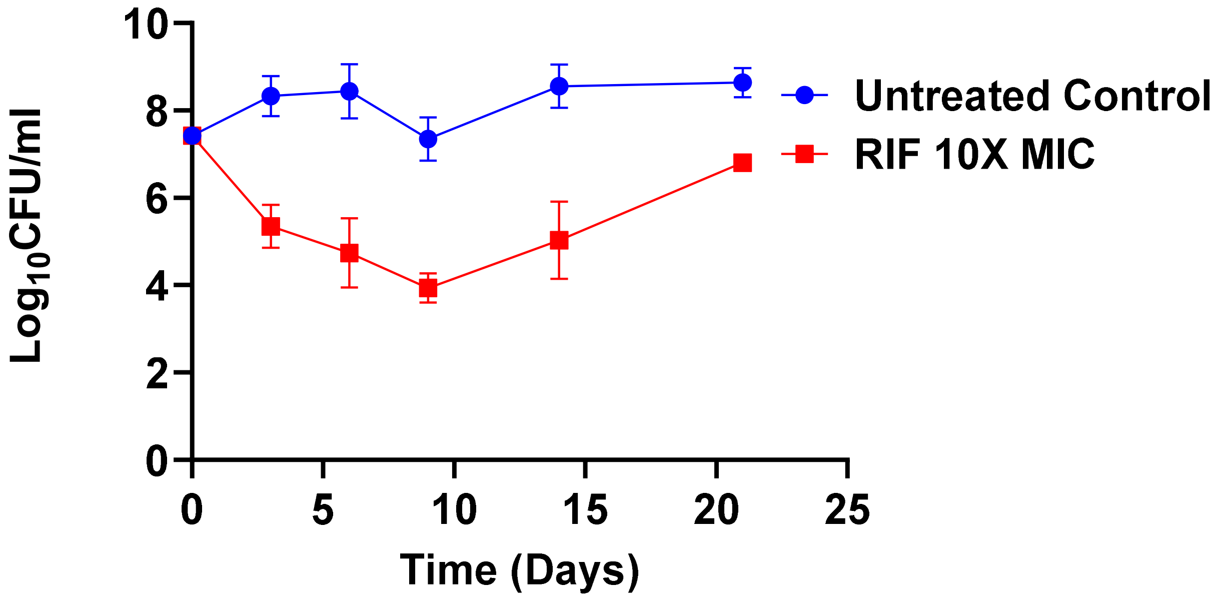


**Figure S3:** Killing kinetics of rifampicin at 10X MIC concentration against *M. tuberculosis*.
